# High resolution profiling of pathways of escape for SARS-CoV-2 spike-binding antibodies

**DOI:** 10.1101/2020.11.16.385278

**Authors:** Meghan E. Garrett, Jared Galloway, Helen Y. Chu, Hannah L. Itell, Caitlin I. Stoddard, Caitlin R. Wolf, Jennifer K. Logue, Dylan McDonald, Frederick A. Matsen, Julie Overbaugh

## Abstract

Defining long-term protective immunity to SARS-CoV-2 is one of the most pressing questions of our time and will require a detailed understanding of potential ways this virus can evolve to escape immune protection. Immune protection will most likely be mediated by antibodies that bind to the viral entry protein, Spike (S). Here we used Phage-DMS, an approach that comprehensively interrogates the effect of all possible mutations on binding to a protein of interest, to define the profile of antibody escape to the SARS-CoV-2 S protein using COVID-19 convalescent plasma. Antibody binding was common in two regions: the fusion peptide and linker region upstream of the heptad repeat region 2. However, escape mutations were variable within these immunodominant regions. There was also individual variation in less commonly targeted epitopes. This study provides a granular view of potential antibody escape pathways and suggests there will be individual variation in antibody-mediated virus evolution.

## INTRODUCTION

The global outbreak of a novel coronavirus, SARS-CoV-2, has claimed over one million lives within just eleven months after the first detected case (https://coronavirus.jhu.edu/map.html), with countless others experiencing long-term health problems after recovering from infection. Vaccines to prevent SARS-CoV-2 spread and treatments to reduce disease severity are currently under rapid development, with many strategies relying on antibody-mediated immunity. The main viral target of interest for vaccines and antibody therapies against SARS-CoV-2 is the coronavirus spike (S) protein, which decorates the surface of the virion and mediates attachment and entry into host cells (Walls et al., 2020). The S protein is comprised of a trimeric assembly of two subunits: S1 and S2, which are proteolytically cleaved at the S1/S2 boundary. The S1 subunit contains an N-terminal domain (NTD) and a receptor binding domain (RBD) within the C-terminal domain (CTD). The S2 subunit contains the fusion peptide (FP) along with two heptad repeat regions (HR1 and HR2), separated by a linker region, responsible for driving viral and host membrane fusion (Walls et al., 2017; Xia et al., 2020). Binding of the S1 protein via the RBD to the human ACE2 receptor is followed by proteolytic cleavage at the S2’ site, which exposes the FP and activates a series of conformational changes resulting in membrane fusion (Belouzard et al., 2009; Fan et al., 2020; Shang et al., 2020).

Neutralizing antibodies targeting the SARS-CoV-2 RBD have been the main focus of vaccine strategies and antibody therapies, as they block virus entry in cell culture (Pinto et al., 2020; Wan et al., 2020; Wang et al., 2020; Wec et al., 2020) and prevent infection or disease in some animal models (Hassan et al., 2020; Rogers et al., 2020; Zost et al., 2020). However, the study of other coronaviruses has illustrated that antibodies elicited by infection can target epitope regions outside of the RBD. For example, a number of neutralizing antibodies directed to regions other than the RBD have been isolated from individuals infected with the closely related viruses SARS-CoV and MERS-CoV (Shanmugaraj et al., 2020). Recent SARS-CoV-2 studies of serum antibodies from COVID-19 patients have led to the identification of neutralization activity directed at linear epitopes found just downstream of the RBD, overlapping the FP, and just upstream of HR2 in the linker region (Li et al., 2020b; Poh et al., 2020). Thus, there may be multiple regions within the SARS-CoV-2 S protein that may shape the viral immune response.

While antibodies to the RBD are a logical initial focus for studies of protective antibodies against SARS-CoV-2, it is not yet known whether they are a correlate of protection for SARS-CoV-2 in humans. A limited understanding of protective immunity is to be expected at the early stage in this new disease, and a broad view is therefore prudent. In this regard, it is important to note that studies from other viruses such as HIV and Ebola have shown that non-neutralizing antibodies are an immune correlate of protection in humans (Gunn et al., 2018; Haynes et al., 2012; Milligan et al., 2015; Saphire et al., 2018). For SARS-CoV-2, antibodies against the S protein likely perform functions other than neutralization given that there is not a direct correlation between the levels of binding to S protein and neutralization titers (Robbiani et al., 2020). Given the incomplete picture we have regarding immunity against SARS-CoV-2, it is important to study the antibody response irrespective of function and/or epitope in order to fill these knowledge gaps.

Given the rapid spread and amplification of SARS-CoV-2 in the population and the high mutation rate of RNA viruses, variants that can evade the immune response are likely to arise. There is strong evidence that immune selection drove the emergence of escape mutants in the S protein of SARS-CoV (Sui et al., 2008) and MERS-CoV (Kleine-Weber et al., 2019), raising the possibility that the same could occur with SARS-CoV-2. Immune selection may be further enhanced by a vaccine if it is not fully protective, making understanding the potential escape pathways of the virus critically important.

There are several recent studies that have examined the effect of select mutations on serum antibody binding to SARS-CoV-2. One study that assessed 82 S protein variants present in circulating SARS-CoV-2 detected a few mutations that resulted in decreased neutralization sensitivity (Li et al., 2020a). However, only a small fraction of the potential mutations that could arise on the S protein were tested. Another study harnessed the power of deep mutational scanning (DMS) to capture a complete picture of the functional consequences of single mutations within the RBD on protein expression, ACE2 binding, and monoclonal antibody binding (Starr et al., 2020). However, no study has yet examined the effect of the polyclonal antibody response on immune escape across the S protein, which is the target in current prophylactic and therapeutic strategies to combat COVID-19.

Previously we developed a comprehensive method of mapping escape mutations for HIV monoclonal antibodies, referred to as Phage-DMS (Garrett et al., 2020). In Phage-DMS, a library of all possible mutations to a protein is generated in peptide fragments, which are expressed by phage. Complexes of antibodies that bind the phage library are immunoprecipitated and sequenced to determine the antibody binding region(s) and the mutations within that epitope region that disrupt binding. Phage-DMS has several advantages: it is high-throughput, allowing comparison amongst a large number of samples in parallel, it can determine pathways of escape for both neutralizing and non-neutralizing antibodies that bind to linear epitopes, and results using this method also correlate well with mutational effects measured in other assays (Garrett et al., 2020). Here, we used Phage-DMS to understand the spectrum of single mutations on the S protein that could reduce antibody binding and thus mediate escape from plasma antibodies found in COVID-19 patients. Using a Phage-DMS library displaying both wildtype and mutant peptides tiling across the S protein, we identified a spectrum of single mutants that were capable of reducing antibody binding and found person-to-person variability in the effect of mutations within immunodominant epitopes. In the arms race between the humoral immune response and SARS-CoV-2, these results allow us to predict pathways of escape and forecast the appearance of escape mutants.

## RESULTS

### Generation of the Spike Phage-DMS library

In order to explore all amino acids that define the epitope of antibodies directed towards the spike (S) protein of SARS-CoV-2, we generated a Phage-DMS library designed to tile across the ectodomain of the S protein from the Wuhan Hu-1 strain (Figure 1). We also included DMS peptides generated in the context of the D614G mutation, as clinical and in vitro evidence suggests that this variant may have increased infectivity as compared to the original Wuhan Hu-1 strain (Korber et al., 2020). We computationally designed sequences coding for peptides 31 amino acids long with the variable amino acid in the central position. To achieve single amino acid resolution of epitope boundaries, peptides were designed to overlap by 30 amino acids. 24,820 unique peptides were designed in total; the peptide library included wildtype peptides that could be used to define the antibody epitope and peptides with all possible mutations to determine those within the defined epitope that disrupt or enhance antibody binding. Two biological replicate libraries of these peptide sequences were cloned as we have done previously (Garrett et al., 2020). Deep sequencing of the final duplicate libraries (Library 1 and Library 2) indicated that each contained a high percentage of all unique sequences (96.0% and 95.9%, respectively) (Figure S1).

**Figure 1.**
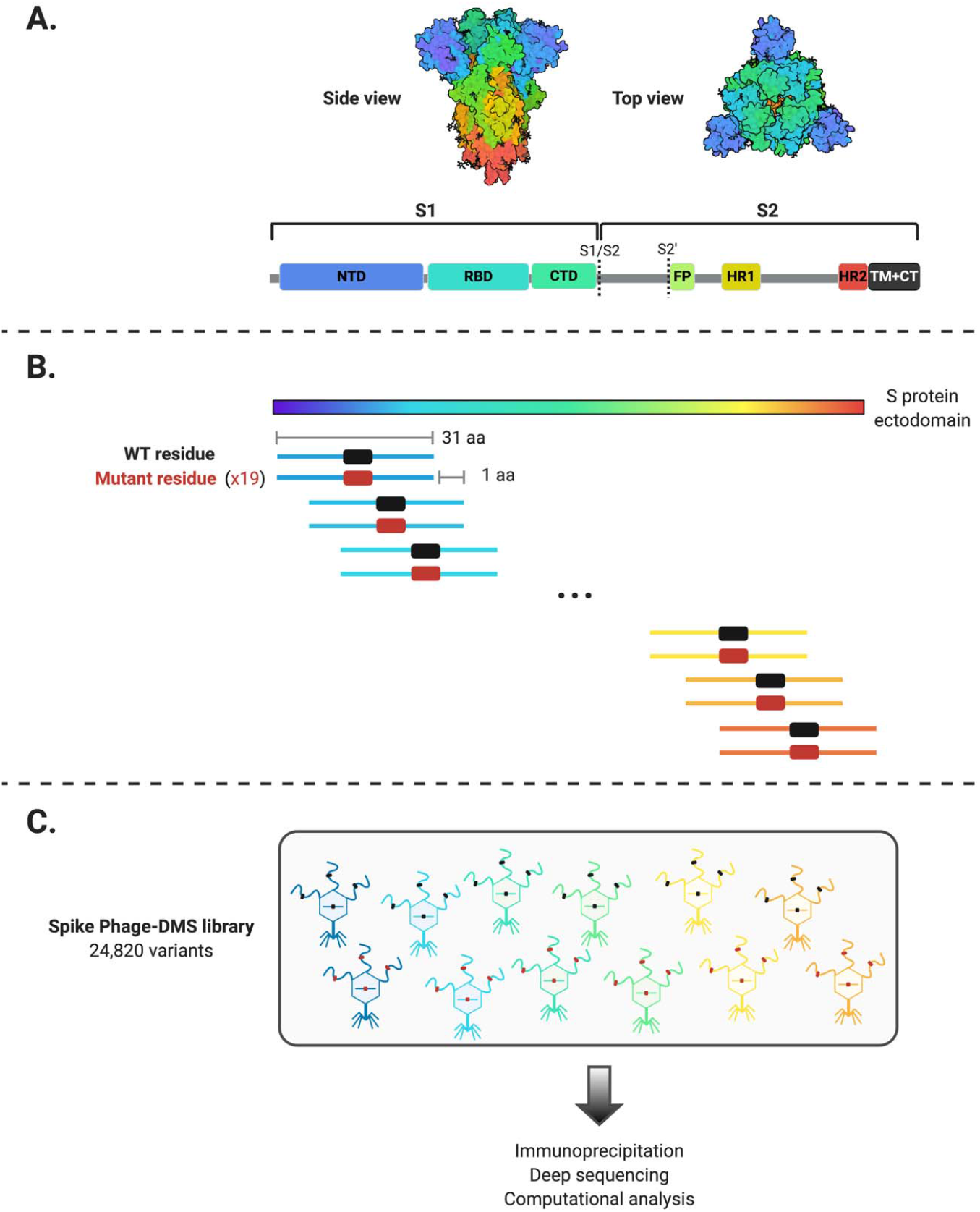
Schematic of the design of the Spike Phage-DMS library. (A) Structure of the S protein and location of important protein domains. Structure was made in BioRender.com (PDB: 6VX**X)**. (B) Sequences were computationally designed to code for peptides 31 amino acids long and to tile stepwise across the Wuhan-Hu-1 SARS-CoV-2 S protein ectodomain by 1 amino acid. There are 20 peptides representing all 20 possible amino acids at the central position, containing either the wild type residue (shown in black) or a mutant residue (shown in red). Within the 31 aa region surrounding the D614G mutation, peptides were also generated with G614 in addition to the 20 amino acid variants at the central position. (C) The designed sequences were cloned into a T7 phage display vector and amplified to create the final S protein Phage-DMS library. This library was then used in downstream immunoprecipitation and deep sequencing experiments with human plasma.

### Enrichment of immunodominant linear epitopes by antibodies from COVID patients

We used the Spike Phage-DMS library to determine the pattern of antibody binding in plasma from a cohort of 18 COVID-19 patients from the HAARVI study in the Seattle area collected between March and May 2020 (Table S1). Most patients had mild symptoms not requiring hospitalization, with the exception of one patient (6) who had moderate symptoms and required supplemental oxygen. Convalescent plasma was collected twice, at approximately day 30 and day 60 post symptom onset (p.s.o.; Table S1).

To define the epitope region targeted by antibodies, we examined the enrichment of wildtype peptides from the library. We first calculated Pearson’s correlation coefficient for the peptide enrichment values from each biological replicate experiment. In general, samples with poor correlation between replicate experiments were those that lacked reproducibly strong antibody binding. The correlation ranged from 0.96 to 0.27, with three patients having no sample from either timepoint above a correlation of 0.5 (Figure S2A). The latter three cases (7, 16, and 17) were excluded from further analyses. Paired samples taken from the same individual at 30 and 60 days p.s.o. had peptide enrichment values that were well correlated, with a median correlation of 0.87, and were significantly better correlated than randomly paired samples (p = 1.2e-09, Wilcoxon rank sum test), which had a median correlation of 0.29 (Figure S2B). The correlation between paired samples also tended to be stronger when the correlation between biological replicates, and therefore antibody binding, was stronger (Figure S2C).

Wildtype peptides from two immunodominant regions, which include the FP and upstream sequences spanning aa 809-834 as well as the linker region and N-terminal portion of the adjacent HR2 domain spanning aa 1140-1168, were the most enriched peptides (Figure 2). Peptide enrichment data is also available in interactive form using the dms-view online tool(Hilton, 2020) at https://github.com/meghangarrett/Spike-Phage-DMS/tree/master/analysis-and-plotting/dms-view. Interestingly, while most patients showed enrichment primarily within the linker region upstream of HR2, patient 5 showed strong enrichment of peptides from an epitope within the HR2 domain itself. The enriched peptides included aa 1167-1191 and this epitope was unique compared to the other 14 patients. We also observed cases where patient plasma enriched peptides from a less common epitope, albeit weakly, at both 30 and 60 days p.s.o.. Plasma from patient 12 enriched peptides within a region of the NTD (aa 255-280), patient 15 enriched peptides within both the RBD (aa 485-500) and the region just downstream of the RBD (aa 540-573), and patient 3 enriched peptides in the region just upstream of the S1/S2 cleavage site (aa 620-644) (Figure 2).

**Figure 2.**
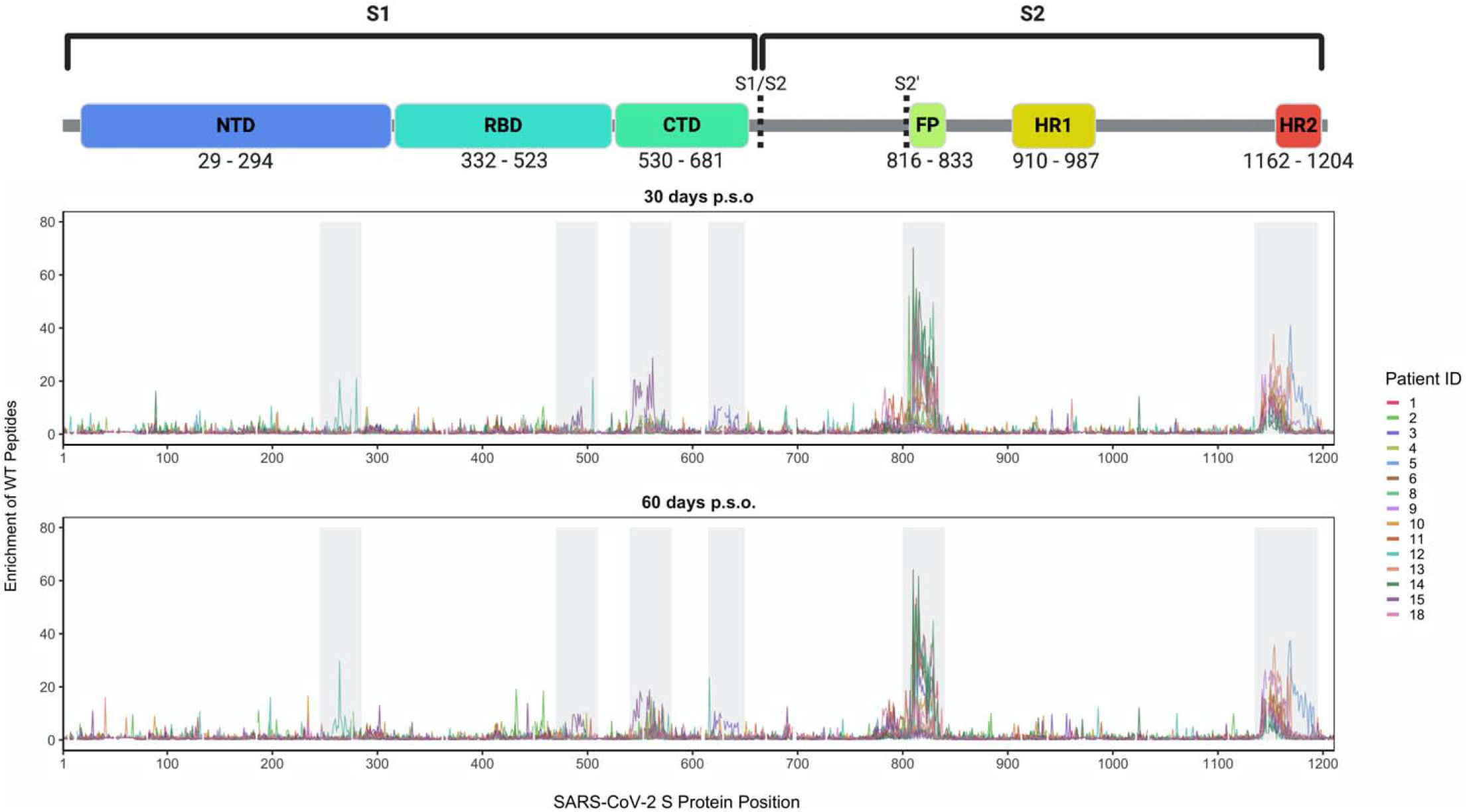
Linear epitopes bound by COVID-19 patient plasma. Lines represent the enrichment of wildtype peptides from the Spike Phage-DMS library from individual plasma samples. Samples from convalescent COVID-19 patient plasma taken at approximately day 30 p.s.o. (top panel) or day 60 p.s.o. (bottom panel) are shown. Lines are colored by patient, with the key to the patient IDs on the right. Grey boxes highlight immunogenic regions where enrichment was detected in at least one individual across timepoints. Peptides that were included in the design, but absent from the phage library (Figure S1), are shown as breaks in the line plots. A schematic of S protein domains is shown above, with locations defined based on numbering used in: https://cov.lanl.gov/components/sequence/COV/annt/annt.comp

### Patient-to-patient variability in mutations that lead to loss of antibody binding

To determine the effect of mutations on antibody binding to S protein epitopes, we compared the relative enrichment of wildtype peptides and mutant peptides within the epitopes defined above. To quantify the effect of each amino acid on binding, we calculated the differential selection of mutant peptides versus wildtype peptides and scaled this value by the strength of binding to the wildtype peptide, as we have previously done for Phage-DMS experiments; this measure is highly correlated with the relative binding of individual mutant peptides by ELISA (Garrett et al., 2020). Plotting the scaled differential selection values for all mutants at each site in a heatmap allows for the visualization of sites where mutations led to a detectable loss of binding. The scaled differential selection data for all patients is also available to view in logo plot form at https://github.com/meghangarrett/Spike-Phage-DMS/tree/master/analysis-and-plotting/dms-view.

We generated heatmaps for four representative samples from COVID-19 patients with strong enrichment. We did this for the two immunodominant regions, FP and linker region/HR2, and included paired day 30 and 60 p.s.o. samples (Figure 3 and 4). We also examined the escape profiles within less common epitopes (Figure S3). While the effect of mutations seemed to be consistent between paired patient samples, between individuals there was more heterogeneity in the specific escape profiles for each region, as follows:

**Figure 3.**
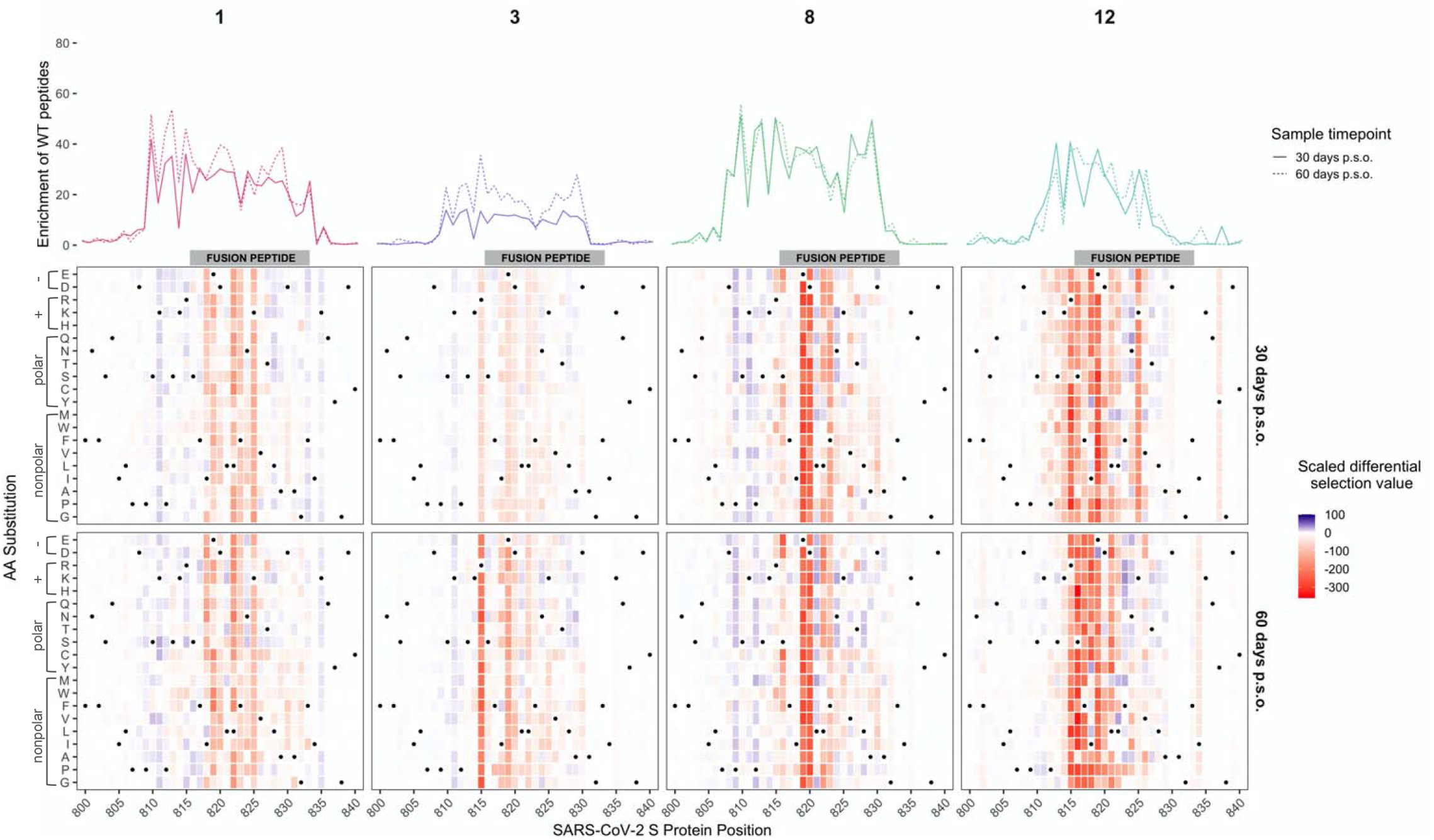
Effect of mutations on binding by COVID-19 patient plasma within the FP region. Heatmaps depicting the effect of all mutations, as measured by scaled differential selection, at each site within the FP epitope for representative COVID-19 patients (numbered at top). Mutations enriched above the wildtype residue are colored blue and mutations depleted as compared to the wildtype residue are colored red. The intensity of the colors reflects the amount of differential selection as indicated to the right. The wildtype residue is indicated with a black dot. Line plots showing the enrichment of wildtype peptides for each patient are shown above, with a solid line for the day 30 p.s.o. patient samples and a dashed line for the day 60 p.s.o. patient samples.

**Figure 4.**
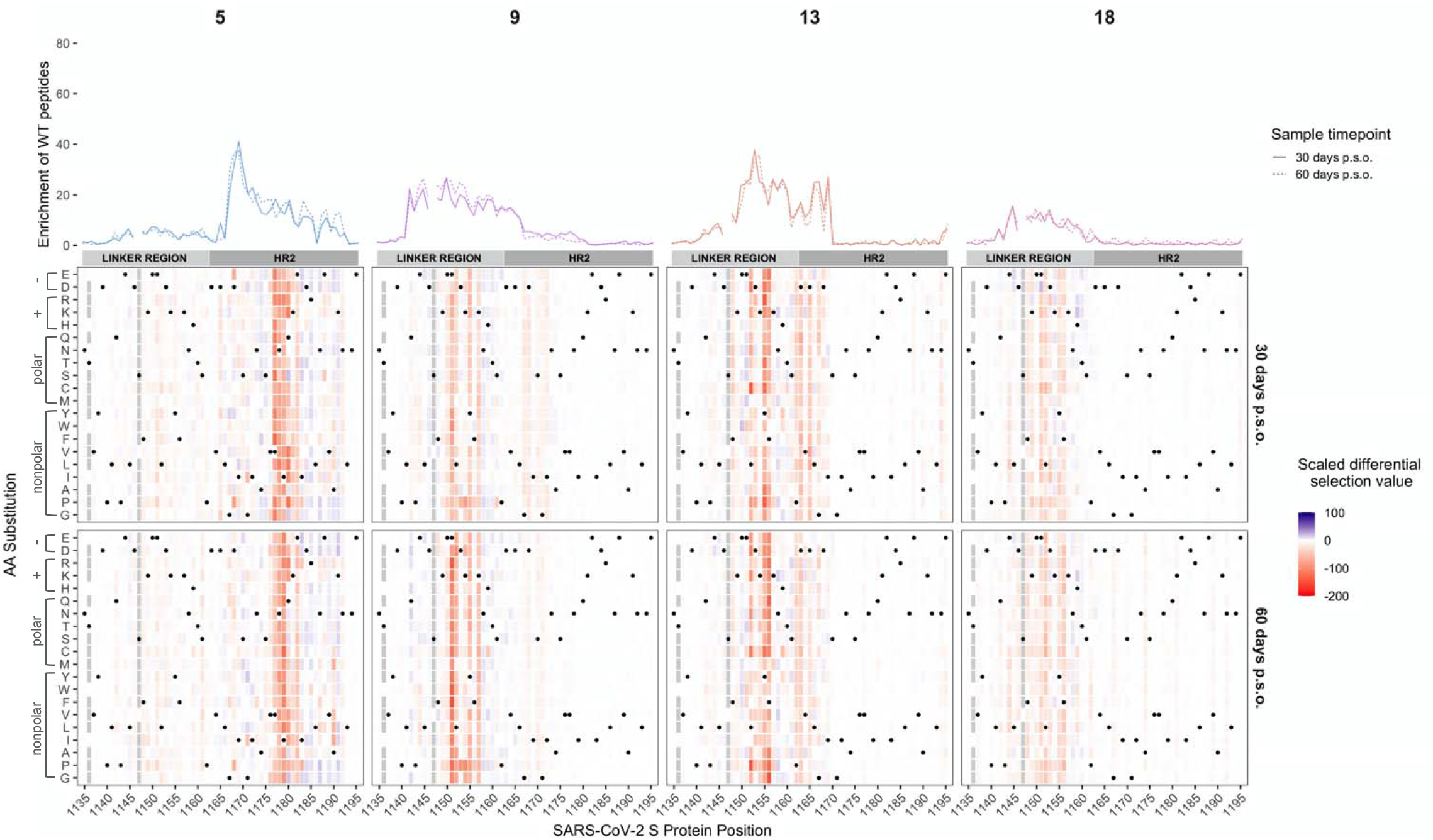
Effect of mutations on binding by COVID-19 patient plasma within the linker/HR2 region. Heatmaps depicting the effect of all mutations, as measured by scaled differential selection, at each site within the linker region/HR2 epitope for representative COVID-19 patients (numbered at top). Mutations enriched above the wildtype residue are colored blue and mutations depleted as compared to the wildtype residue are colored red. The intensity of the colors reflects the amount of differential selection as indicated to the right. The wildtype residue is indicated with a black dot. Line plots showing the enrichment of wildtype peptides for each patient are shown above, with a solid line for the day 30 p.s.o. patient samples and a dashed line for the day 60 p.s.o. patient samples. Peptides missing from the library are shown as grey boxes in the heatmaps and as breaks in the line plots.

#### FP region

Within the FP region, sites that led to reduced binding clustered primarily within the 5’ region of the FP itself, which spans from sites S816 to F833 (Figure 3A). Some sites were similarly sensitive between patients. For example, mutations at site 819 and 820 disrupted antibody binding to some extent in all of the patients shown and were the dominant escape positions for patient 8. In contrast, mutations in the adjacent site 818 appeared to contribute to the epitope of some patient antibodies (1, 3, 12) but not others (8). One interesting site that variably contributed to antibody binding is R815, which is upstream of the FP cleavage site and thus would not be predicted to be present on the postcleavage form of S2. Antibodies from patient 3 and 12 both showed reduced binding to peptides with mutations at this site, indeed, this was the dominant escape mutation for patient 3. Substitutions at this same amino acid position had a more modest impact on binding and only a few amino acids led to disruption for patient 8. In the case of patient 1, several mutations at 815 had a positive differential selection value, suggesting they enhance binding.

Not all mutations at each site were equally disruptive to epitope binding, suggesting specific mutations rather than removal of the wildtype amino acid per se may be more important for escape. For example, addition of negatively charged amino acids at site S816 led to loss of binding for patient 8, whereas addition of small amino acids such as alanine or glycine at the same site had little to no effect. We noted some cases where a mutation had a positive differential selection value and thus presumably bound more strongly than the wildtype SARS-CoV-2 sequence at that position. We aligned the FP sequences for SARS-CoV-2 and human endemic coronaviruses and found that some mutations selected above wildtype were residues present in FP sequences from other CoVs (Figure S3). For patient 12, a few mutants at site 818 including I818L had positive differential selection values, and two coronaviruses (HKU1 and NL63) have a leucine at position 818. In another instance, the N824S mutation had a positive scaled differential selection value for all four samples taken 60 days p.s.o., and we saw that NL63 and 229E both have a serine at position 824.

#### Linker/HR2 region

Within the region surrounding HR2, we observed that most patients targeted an epitope spanning aa 1123-1162 within the linker region just upstream of the HR2 domain, with contribution of residues within the N terminus of the HR2 domain in some cases, while one patient targeted an epitope within the HR2 domain itself (Figure 4). In addition to this individual variation in the epitope boundaries, the mutations at different sites exhibited marked variability between patients, as seen for the FP epitope. An example of this variability can be seen among patient antibodies targeting distinct epitopes within the linker region. In this epitope, patient 13 exhibited sensitivity to mutations spanning 1162 to 1167, while this was not seen in the other patients. There were also some cases where there were similarities in escape profiles for patient antibodies targeting the same region. For example, there was evidence that amino acid changes at sites 1151 and 1152 disrupted antibody binding in several patients. However, while mutations at site 1151 disrupted antibody binding for patients 9 and 18, they did not for the antibodies in patient 13.

In patient 5, where there was a distinct epitope within the HR2 domain, we observed unique sensitivity to mutations in sites 1176 through 1182 not found in any other patient. Like many other plasma epitope profiles, mutations to the same site had a variable effect. For example, at site 1180 some mutations improved binding (ex: Q1180G), some reduced binding (ex: Q1180L), while some had little to no effect (ex: Q1180A). There was no overlap in the sites that showed reduced binding in patient 5 and those that were negatively selected in patients 9, 13, or 18, again suggesting that the linker region and HR2 domain epitopes are distinct epitopes that have distinct pathways of escape.

#### Other regions

There was relatively less differential selection at regions outside of the immunodominant epitopes, but the profiles nonetheless provided evidence for some specific mutations that disrupt binding in epitopes in the NTD, RBD, and CTD just downstream of the RBD (Figure S4). For example, within the NTD, positions spanning 264-269 in patient 12 disrupted binding, most notably mutations from A to select amino acids (W, V and I) at site 264. The escape profiles for patient 15 suggests that changes at positions 491-495 disrupt binding in the RBD epitope, while mutations at sites 550-553 and 558 disrupted binding within the CTD epitope for this patient. For patient 3, the major effect on binding within the CTD was at positions 628-634, most notably at amino acids 628 and 633. Although negative scaled differential selection values can be seen at various other amino acid positions, in many of these cases the effects were weak and/or inconsistent across time points. Identifying cases of stronger enrichment in these regions will be necessary in order to better define the full spectrum of escape mutations at these regions.

### Most common circulating SARS-CoV-2 variants, including D614G, are not predicted to escape antibody binding

One question that arises from the escape profiles described above is whether the mutations that disrupt antibody binding identified here correlate with the global evolution of the SARS-CoV-2 virus to-date. Using variant frequencies present in sequences from GISAID and reported at the https://cov.lanl.gov website, we compared the naturally occurring diversity of every site to the effect on antibody binding as found by Phage-DMS. For each patient and at each site, we averaged the scaled differential selection value for all mutants and then plotted this against the mutational entropy, which is a measure of amino acid diversity (Figure 5A). Larger mutational entropy values indicate more global diversity at that site, and sites with a mutational entropy value of above 0.02 are flagged as sites of interest by the LANL database. We found that the majority of sites that led to loss of antibody binding when mutated were not present at a high frequency in nature. Sites within the immunodominant epitopes did not generally have high mutational entropy, although for patient 13 there were a few sites within the HR2 region that were both present at high frequency in nature (mutational entropy > 0.02) and led to loss of antibody binding when mutated.

**Figure 5.**
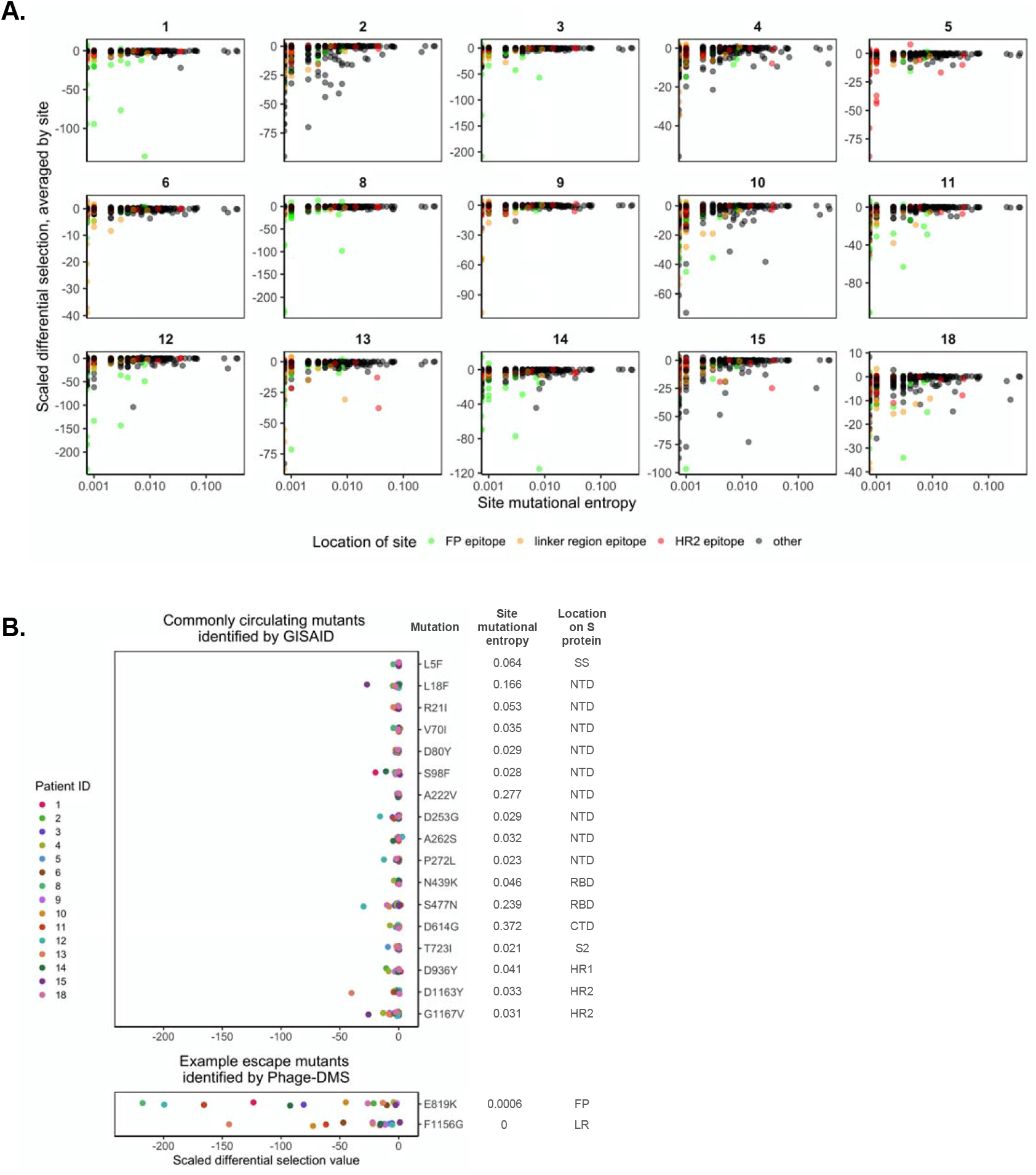
Predicted effects of commonly circulating S protein variants on antibody escape. (A) Scatterplot comparing the effect of mutations on patient plasma antibody binding and the frequency of all circulating S protein variants. The mutational entropy of every circulating protein variant, as reported at the https://cov.lanl.gov website and based on GISAID global sequencing, is plotted on the x-axis. The average of the scaled differential selection values for all mutants at each site is plotted on the y-axis. Patient ID’s are indicated on the top. Each site is colored by its location, as indicated on the bottom. (B) Effect of mutant peptides representing commonly circulating S protein variants on binding to COVID-19 patient plasma. We selected sites with a mutational entropy of greater than 0.02, as this is the cutoff used by LANL to determine sites of interest. On the top are the 17 sites of high mutational entropy and on the bottom are two selected sites that were noted as sites of antibody escape within immunodominant epitopes by Phage-DMS. On the right are the mutations examined, named according to the wildtype aa, followed by the site number, followed by the mutant aa of interest. Mutations chosen at sites of high mutational entropy represent the most common variant found in nature. The scaled differential values found by Phage-DMS for each mutant peptide are shown as dots and are colored by patient as indicated to the left. Data is from samples taken day 60 p.s.o.. SS = signal sequence, S2 = N-terminal region of S2, LR = linker region.

Focusing on circulating variants that are rising in abundance in the S protein ectodomain, we next examined the effect of mutations at these sites as determined by Phage-DMS. The 17 selected sites with a mutational entropy of greater than 0.02 were located across the S1 and S2 proteins and mostly were not found within the major immunodominant regions identified here, with the exception of two mutations that appear within the HR2 domain. We chose the most commonly found natural variant present at each site of high mutational entropy and examined the scaled differential selection value of that variant for all 60 day patient samples. In general, the variants found in the global population did not show reduced binding to patient plasma antibodies, with perhaps a modest effect for a few mutations with individual patient samples (Figure 5B). This contrasted with the strong negative selection observed in the Phage-DMS screen for mutations within the two immunodominant epitopes at position 819 and 1156, two positions where there are not variants rising in abundance in nature. However, the possibility remains that sites of high mutational entropy could exist within conformational epitopes, which are not generally displayed in phage libraries.

### Evidence for patient-specific epistatic effects of D614G

We were particularly interested in the S protein mutant D614G, a variant that has emerged as the dominant circulating strain of SARS-CoV-2. Because this library also contains peptides made in the background of the D614G strain, we were able to test whether this mutation could, through epistasis, interact with mutations introduced at other sites on the same peptide and together modify the ability of antibodies to bind. For all mutant peptides that tiled across site 614, the enrichment of each peptide made in the original Wuhan Hu-1 strain background (D614) was compared against the enrichment of the corresponding peptide made in the background of the G614 mutation. We found that the presence of the D614G mutation significantly reduced the ability of antibodies from patient 10 to bind to mutant peptides in this region, with a moderate effect size (Figure 6, Wilcoxon paired signed-rank test). In 8 out of the other 14 patients there was a statistically significant difference in the binding between mutant peptides with and without the D614G mutation, but the effect sizes were all small. Occasionally, for example in the case of patient 4, a mutation at one position (A609W) showed stronger enrichment in the context of G614 as compared to D614. Conversely, in the case of patient 1, one mutation (A609D) was more enriched when accompanied by D614 as compared to G614. In other cases, such as with patient 2, there was some selective enrichment in both contexts.

**Figure 6.**
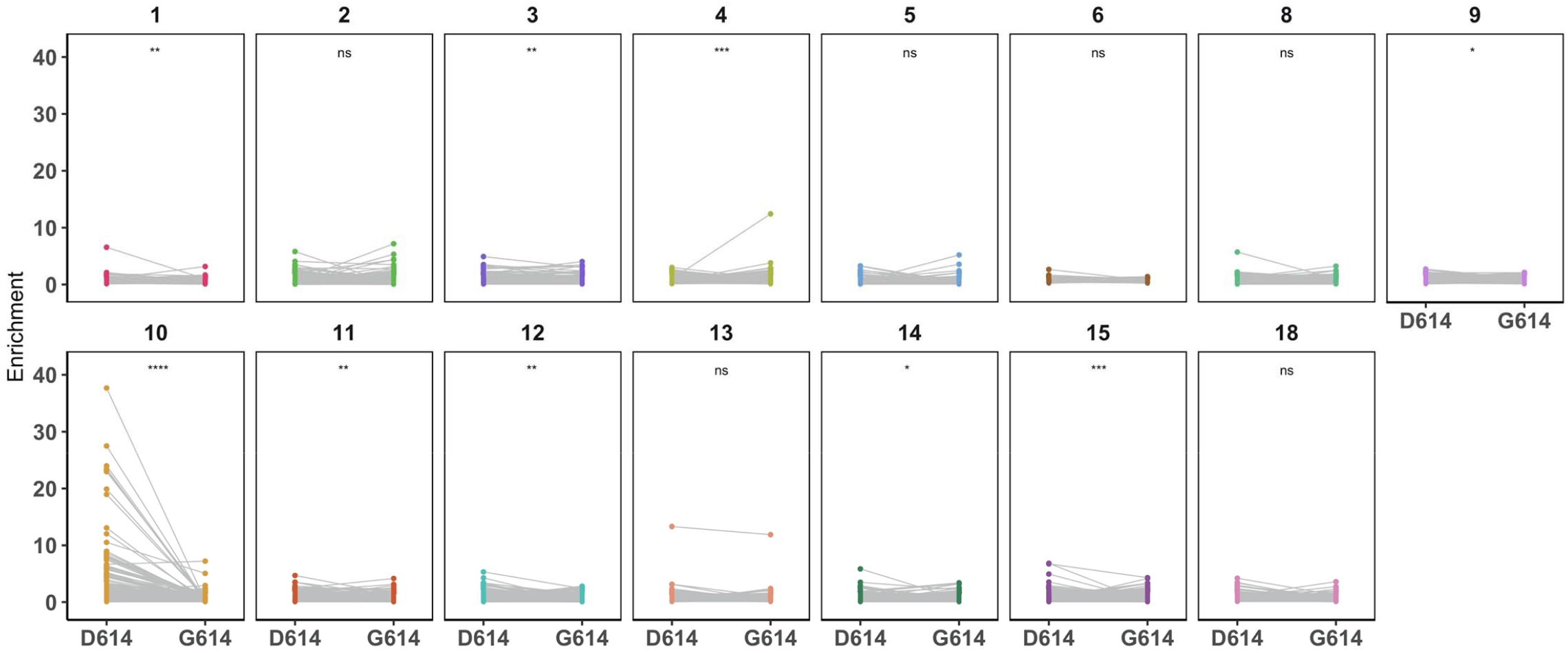
Epistatic effects of D614G mutation on antibody binding. Enrichment values for paired mutant peptides made in either the wildtype Wuhan Hu-1 strain (on the left, D614) or D614G background (on the right, G614) for each patient (numbered at top). All mutant peptides that contained site 614 were included in this analysis (spanning aa 599-629). Data is from samples taken day 60 p.s.o.. Wilcoxon paired signed-rank test was performed (n = 380 paired mutant peptides). The effect size for all patient samples was small (Wilcoxon *r* < 0.3) except for patient 10, whose antibodies exhibited a moderate effect (Wilcoxon *r* = 0.46).

## DISCUSSION

In this study, we tested all possible mutations on the S protein to provide a map of escape pathways within immunodominant linear epitopes targeted by the plasma of convalescent COVID-19 patients. The responses within an individual were consistent over time, but there were many unique pathways of escape that differed between individuals, even within the same epitope region. These findings suggest that the pattern of virus evolution within the growing pandemic is not likely to be driven by a single antibody escape mutation, which may explain the lack of emergence of circulating strains with mutations that disrupt antibody binding identified here. Thus, the responses to a SARS-CoV-2 S protein vaccine immunogen are not likely to be uniform, nor will the pathways of escape.

While others have defined the linear epitopes bound by antibodies within COVID-19 patient serum (Jiang et al., 2020; Li et al., 2020b; Poh et al., 2020; Shrock et al., 2020; Yi et al., 2020; Zamecnik et al., 2020), there have been limited studies to determine what mutations within these regions could abrogate antibody binding. One study that tested 82 circulating SARS-CoV-2 variants showed that the few single mutations that did result in reduced neutralization sensitivity were unique to each serum sample (Li et al., 2020a), and our data supports and extends the idea that antibody binding sites and escape mutants vary greatly from person to person. The lack of a singular escape signature within each immunodominant epitope also implies that a diverse repertoire of antibodies targets these regions. This provides a functional context for results of studies of SARS-CoV-2 S protein antibodies sequences from COVID-19 patients, which showed that no single clones dominate the antibody response; rather, a diverse collection of variable heavy and light chain genes are used in different individuals (Brouwer et al., 2020; Seydoux et al., 2020).

Our results also demonstrate the power of interrogating the role of every possible amino acid at every site on the S protein. Mapping COVID-19 patient serum epitopes by alanine scanning has helped identify sites of antibody binding by removing important side chains interactions (Shrock et al., 2020), but studies with other viruses have shown that escape can be mediated by mutations at sites not directly in contact with the antibody via introduction of nearby charged or bulky amino acids (Dingens et al., 2019; Doud et al., 2017; Patel et al., 2019). This concept was evident in this study, for example within the FP epitope for patient 8, where addition of negatively charged amino acids led to escape at site S816 but addition of an alanine did not. We were also able to examine the effects of mutations in the context of the common circulating 614 variant, which we found did not itself drive escape from antibody binding. However, our data suggests that mutation D614G does potentiate escape mutations at other positions in at least one patient, highlighting the power of examining combinations of mutations to better understand the increasing global dominance of the D614G variant.

The FP and linker region/HR2 immunodominant epitopes profiled in this study offer intriguing alternative targets for vaccine design or immunotherapy that could complement efforts focused on the RBD. S2 in general, and the FP in particular, is highly conserved among coronaviruses, indicating the strong purifying evolution acting in this region due to functional constraints. The FP is a target of neutralizing antibodies in SARS-CoV-2 patients (Li et al., 2020b; Poh et al., 2020), and these antibodies could act by preventing protease-mediated cleavage at the S2’ site. Interestingly, we saw evidence of antibody binding to sites upstream of the S2’ site in some patients (3 and 12), but not others (1 and 8), potentially indicating that antibodies to the FP could bind during different stages of cell entry, before or after S2’ cleavage. HR2 and the upstream linker region are both targets of neutralizing antibodies in SARS-CoV-2 patients (Li et al., 2020b), and a neutralizing antibody raised against the linker region from murine SARS-CoV infection has been shown to cross-react with SARS-CoV-2 (Lip et al., 2006; Zheng et al., 2020). Antibodies targeting the heptad repeat regions, which undergo large conformational transformations in order to facilitate membrane fusion, could neutralize by binding to the fusion intermediate state. These more conserved targets may be important for designing optimal and durable vaccines, given that RBD has higher mutational entropy, increasing the potential for immune escape from vaccine-induced antibodies. The mutational flexibility of the RBD is supported by *in vitro* studies with RBD-targeting neutralizing antibodies, which found that escape mutants in the RBD of SARS-CoV-2 were rapidly selected (Baum et al., 2020).

There are several important caveats to the results we obtained with Phage-DMS libraries. In general, phage libraries display linear peptides and therefore miss antibodies that bind to complex conformational epitopes, or at best provide a partial view of a linear portion of such epitopes. Additionally, the peptide libraries are amplified in bacteria and therefore lack glycans or other post-translational modifications. While these Phage-DMS-derived escape maps define mutations that could lead to loss of antibody binding, it is unknown whether the virus would tolerate mutations at those sites. In fact, a study examining the effect of mutations within the SARS-CoV-2 RBD demonstrated that many led to poor protein expression or loss of function (Starr et al., 2020). Thus, for the antibody targeting regions described here, which are in more conserved functional domains, the escape mutations should be further examined in the context of their mutational tolerance.

These studies have defined common and variable escape mutations across 18 COVID-19 patients that will be useful for viral surveillance, particularly as SARS-CoV-2 S protein-based vaccines are introduced into the population. In addition, the Spike Phage-DMS library developed here could be useful for examining larger cohorts, potentially including those with variable clinical outcomes and individuals of variable ages, to define whether mutations that disrupt antibody binding vary in a systematic way across populations and whether this is correlated with clinical outcome or risk of reinfection.

## ACKNOWLEDGMENTS

We gratefully acknowledge Kevin Sung for helpful discussion and Katherine H. D. Crawford for Python code used to generate the oligonucleotide library sequences. We thank Sarah K. Hilton, John Huddleston, and Jesse D. Bloom for developing and demonstrating the dms-view online tool. We thank the Fred Hutch Genomics core facility, and in particular Cassie Sather, for assistance with sequencing. This work was funded by NIH grants AI138709 (PI Overbaugh), R01 AI146028 and U19 AI128914 (PI Matsen). Julie Overbaugh received support as the Endowed Chair for Graduate Education (FHCRC). The research of Frederick Matsen was supported in part by a Faculty Scholar grant from the Howard Hughes Medical Institute and the Simons Foundation.

## CONTRIBUTIONS

JO conceived the project; MEG and JO led the design of the study; HYC led the HAARVI study, with CRW, JKL, and DM involved in sample collection. MEG and HLI performed experiments, with CIS performing sample processing and organization; MEG and JG performed computational and data analyses, with FAM advising. MEG and JO wrote the paper with input from all authors.

## STAR METHODS

### KEY RESOURCES TABLE

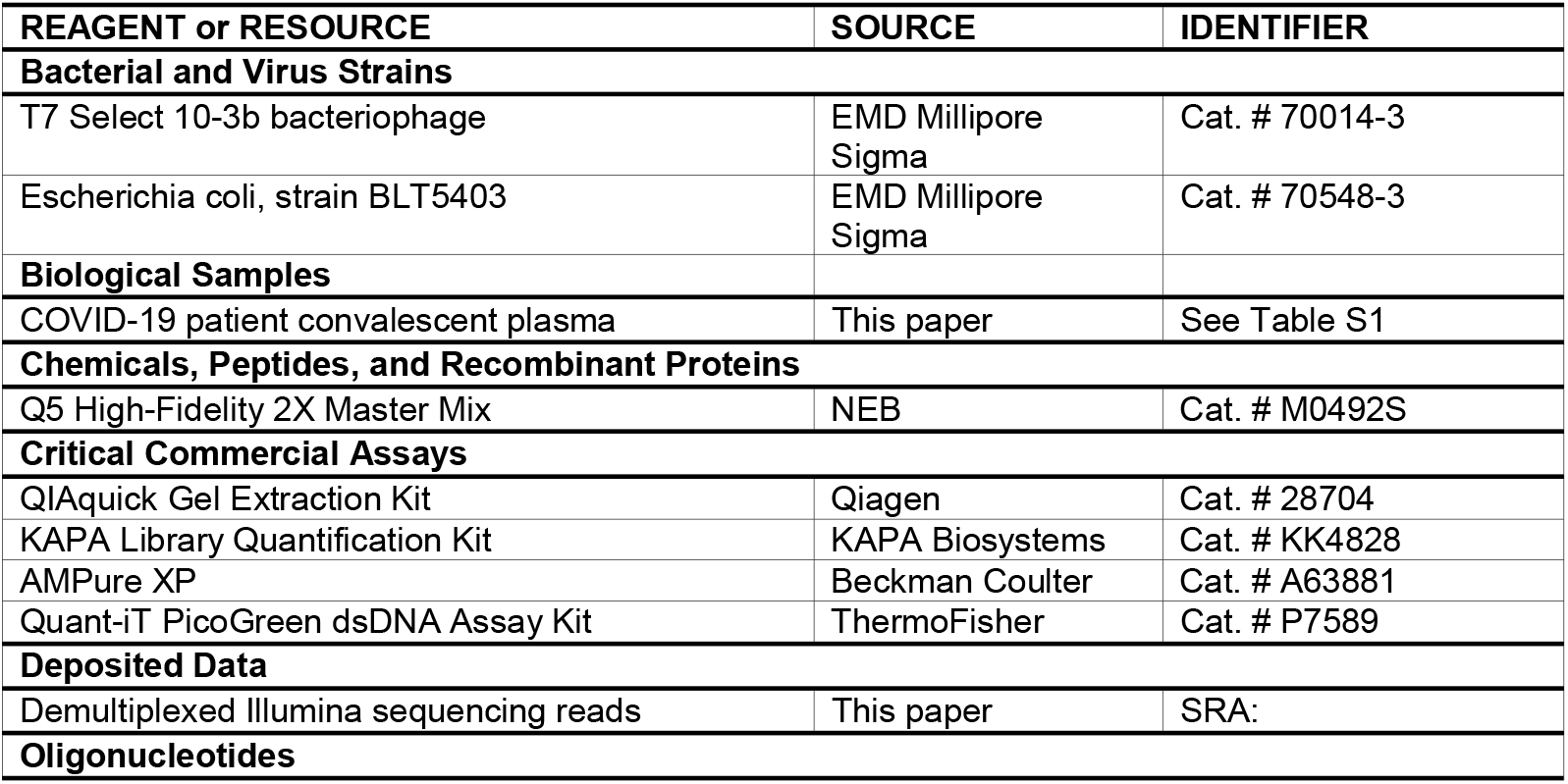
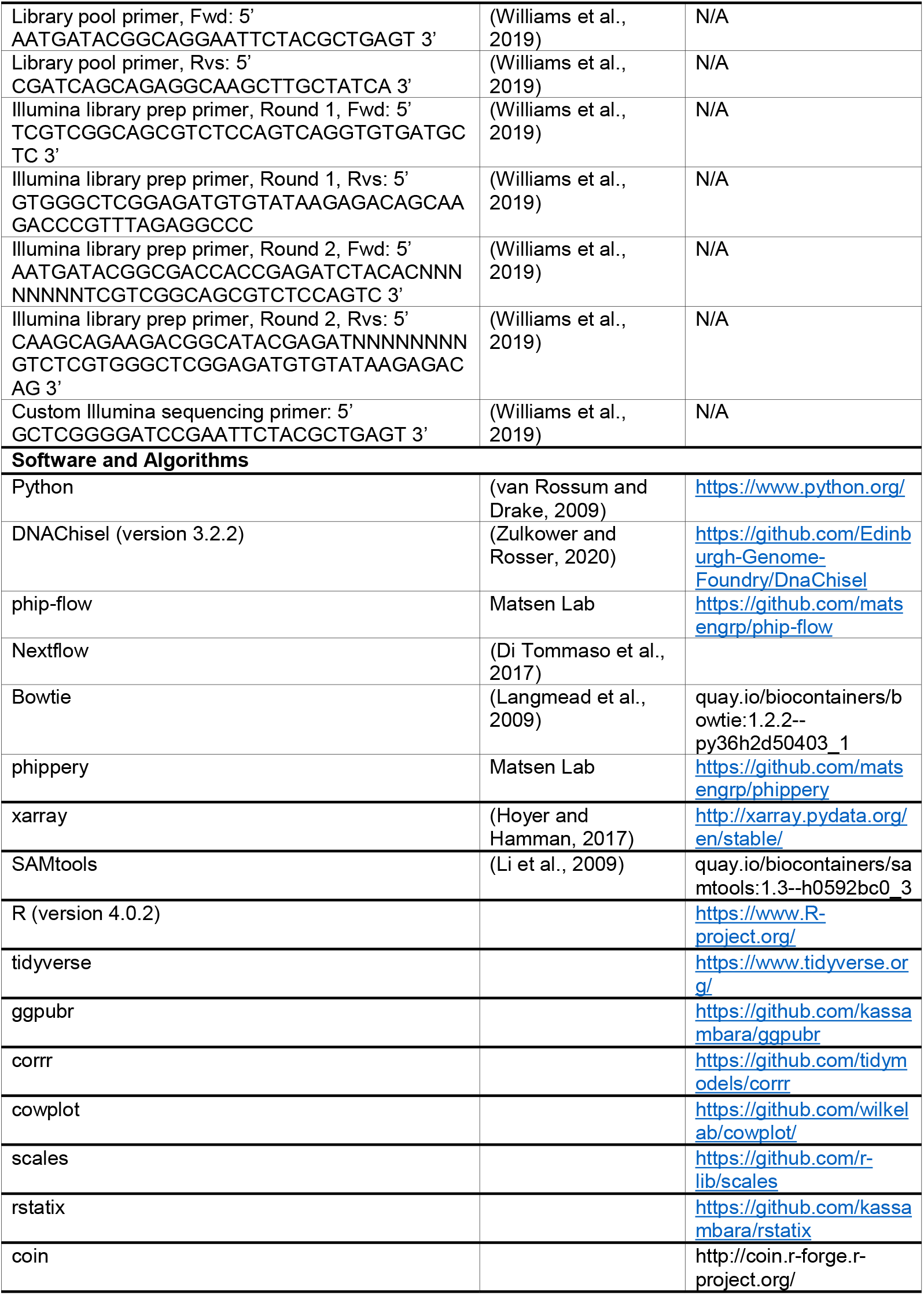
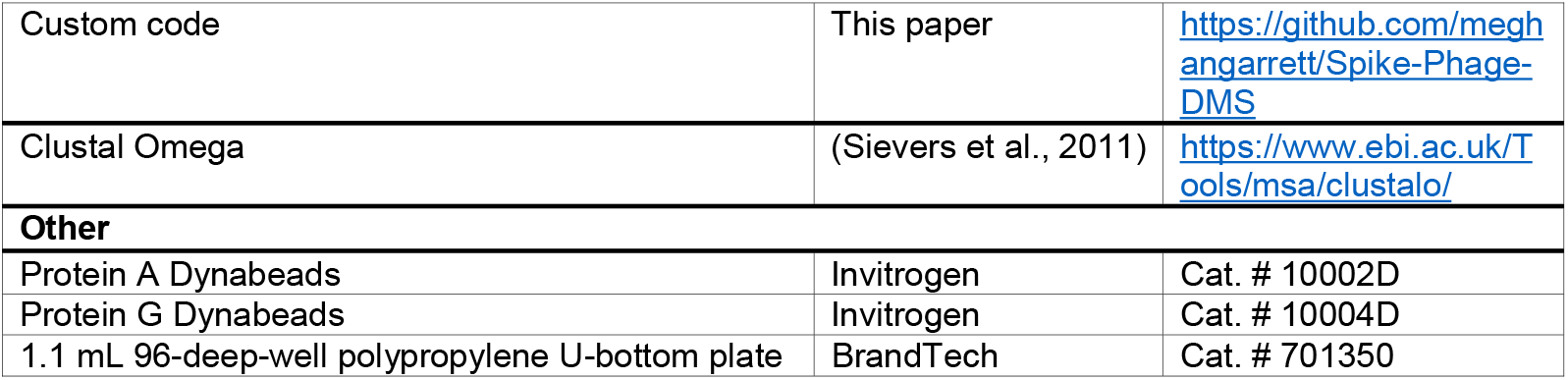

### RESOURCE AVAILABILITY

#### Lead Contact and Materials Availability

Further information and request for reagents may be directed to the corresponding author Julie Overbaugh.

#### Data and Code Availability

Sequencing data has been deposited to NCBI and are accessible under BioProject # XXX. The Nextflow pipeline, used to align all sample reads to the reference library, is available at https://github.com/matsengrp/phip-flow. A custom python package used for all downstream sample curation and analysis is available at https://github.com/matsengrp/phippery. All custom code and input files used to (1) generate oligonucleotide sequences for the S protein Phage-DMS library, (2) run the alignment pipeline, (3) analyze sequencing data for the experiments in this paper, and (4) to generate figures have been deposited at https://github.com/meghangarrett/Spike-Phage-DMS. Data is available to explore using the dms-view website, hosted here: https://github.com/meghangarrett/Spike-Phage-DMS/tree/master/analysis-and-plotting/dms-view.

### EXPERIMENTAL MODEL AND SUBJECT DETAILS

#### Human Subjects

Plasma samples were taken from non-hospitalized COVID-19 patients enrolled in the Hospitalized or Ambulatory Adults with Respiratory Viral Infections (HAARVI) study at the University of Washington. Prior to study initiation, the following institutional human subjects review committee approved the protocol: University of Washington IRB (Seattle, Washington, USA) and concurrent approvals were obtained from the Fred Hutchinson Cancer Research Center for the current study. All plasma samples were heat-inactivated at 56 °C for 1 hour before storage and use. Plasma samples were spun in a centrifuge for 10 min at 1,000 x g in order to clarify the supernatant before use.

### METHOD DETAILS

#### Design and generation of the Spike Phage-DMS library

To create a Phage-DMS library for the S protein of SARS-CoV-2 we used the sequence from the Wuhan Hu-1 strain (Genbank: MN908947). Only the ectodomain of the S protein was included (aa 1-1211), excluding the transmembrane and cytoplasmic domains. Additionally, in the region of the D614G variant of the Wuhan Hu-1 strain, we designed DMS peptides spanning positions 599 through 619 that also include the D614G mutation. Sequences were optimized for uniform GC content (to reduce later biases during PCR amplification) and codon usage for expression in *E coli.* GC and codon optimization was done using the Python package DNAChisel (version 3.2.2), aiming for GC content of between 0.4 and 0.6 within a window of 100 nucleotides. After optimizing the sequences, the two subunits of the S protein (S1 and S2) were then treated as separate proteins. Sequence coding for a glycine-serine linker ([G_4_S]_3_) was added to the beginning and end of the sequence of each protein in order to ensure that the first amino acid of the protein was located in the central position of the peptide. We then generated sequences coding for peptides 31 amino acids long, tiling by 30 amino acids, and containing a single variable residue at the central position of the peptide. This resulted in 20 peptide sequences containing all possible mutations at each position along the proteins, with only one amino acid shift between sequential peptides. Each sequence additionally had 5’ and 3’ adaptor sequences added to facilitate amplification and cloning (5’: AGGAATTCTACGCTGAGT, 3’: TGATAGCAAGCTTGCC). After removing duplicate sequences, 24,820 unique sequences were synthesized by Twist Bioscience as an oligonucleotide pool. Two biological duplicate libraries were generated by independently cloning the sequences into a T7 phage vector and then amplifying the phage, as we have done previously (Garrett et al., 2020). Peptides are numbered by the corresponding S protein location of the amino acid in the central position of the peptide.

Code used to optimize and generate peptide sequences for the S protein Phage-DMS library can be found at https://github.com/meghangarrett/Spike-Phage-DMS.

#### Immunoprecipitation of human plasma with Phage-DMS library

Immunoprecipitation of phage-antibody complexes was performed as previously described (Garrett et al., 2020; Mohan et al., 2018). Briefly, deep 96-well plates were blocked with 3% BSA in Tris-buffered saline with 0.01% Tween (TBST) overnight at 4°C. The phage library was diluted to a concentration representing 200,000 pfu/mL per unique peptide and 1 mL of the diluted phage was added to each well. We assume that plasma IgG concentrations are about 10 ug/uL (Mabuka et al., 2012) and added 10 ug of each sample to the appropriate wells. For every experiment, samples are run in technical duplicate on the same plate. Plates are sealed and rocked at 4°C for 18-20 hours. To immunoprecipitate the phage-antibody complexes we added 40uL of a 1:1 mixture of Protein A and Protein G Dynabeads to each well and incubated the samples for 4 hours at 4°C while rocking. Dynabeads were magnetically separated and then beads were washed 3x with 400 uL wash buffer (150 mM NaCl, 50 mM Tris-HCl, 0.1% [vol/vol] NP-40, pH 7.5). We resuspended beads in 40 uL of water and then lysed bound phage at 95°C for 10 minutes. Additionally, we lysed 10-20 million phage from the diluted input library to determine the distribution of phage in the starting library. Lysed samples were stored at −20°C before preparing for Illumina sequencing.

#### Illumina library preparation and deep sequencing

Lysed phage DNA from each sample was amplified and readied for Illumina deep sequencing by performing two rounds of PCR, as previously described (Garrett et al., 2020). Each PCR reaction was performed using Q5 High-Fidelity 2X Master Mix. For the first round of PCR, 10uL of lysed phage was used as the template in a 25 uL reaction. For the second round of PCR, 2 uL of the round 1 PCR product was then used as the template in a 50 uL reaction, with primers that add dual indexing sequences on either side of the insert. PCR products were then cleaned using AMPure XP beads and eluted in 50 uL water. DNA concentrations were quantified via Quant-iT PicoGreen dsDNA Assay Kit. Equimolar amounts of DNA from the samples, along with 10X the amount of the input library samples, was pooled, gel purified, and the final library was quantified using the KAPA Library Quantification Kit. Pools were sequenced on an Illumina MiSeq with 1×125 bp single end reads using a custom sequencing primer.

#### Multiple sequence alignment

Alignment of FP amino acid sequences from SARS-CoV-2, OC43, HKU1, NL63, and 229E was done using Clustal Omega. GenBank sequences used are as follows: YP_009724390.1, YP_009555241.1, YP_173238.1, YP_003767.1, and NP_073551.1.

#### Graphical illustrations

Graphical abstract, Figure 1, and the diagram of S protein domains in Figure 2 were made in BioRender.com

### QUANTIFICATION AND STATISTICAL ANALYSIS

#### Demultiplexing and alignment of Illumina reads

Demulitplexing and fastq file generation were performed by the Fred Hutch Genomics Core using Illumina MiSeq Reporter software. Demultiplexed sample reads were aligned to the reference library in parallel using a Nextflow data processing pipeline. The pipeline builds a Bowtie index from the peptide metadata by converting the metadata to fasta format and feeds it into the bowtie-build command. The low-quality end of the reads is trimmed to 93bp in order to match the reference lengths before performing end-to-end alignment and allowing for 0 mismatches. For each sample, we quantified the abundance of each peptide by using samtools-idxstats to count the number of reads mapped to each specific peptide in the reference library. The peptide counts were merged into an enrichment matrix organized by unique identifiers for each peptide and sample. The metadata tables were tied with the enrichment matrix into an xarray dataset using shared coordinate dimensions of the unique sample and peptide identifiers. We used this dataset organization as the starting point for all downstream sample curation and analysis.

#### Calculating enrichment and scaled differential selection of peptides

Enrichment and scaled differential selection were calculated as described previously (Garrett et al., 2020). To calculate enrichment, each peptide’s pseudocount frequency was divided by a respective library pseudocount. Each pseudo count was defined as the raw count plus the ratio of the sum of each sample and library count, with a minimum value of P=1. Differential selection of mutant amino acids was then calculated as the log-fold change between each mutant peptide and the wildtype peptide at a locus. By definition, the differential selection of a wildtype amino acid is always 0. The scaled differential selection of a mutant amino acid was then calculated by multiplying the differential selection of a mutant amino acid by the enrichment of the wildtype peptide at that position.

#### Data curation

All samples were screened at least once with each Spike Phage-DMS Library 1 and Library 2, but occasionally were screened multiple times with each library. To ensure we used the same amount of data with each sample, we examined the Pearson’s correlation value of enrichment values for all peptides between all biological replicate experiments and chose the data from the best two correlated experiments for inclusion in the analyses presented here. If there was not at least one timepoint where the sample had a correlation of at least 0.5, then all samples from that patient were excluded from enrichment and scaled differential selection analyses. Peptides that were never sequenced in any experiment were removed from the analyses, and all enrichment and scaled differential selection values shown are the average of two biological replicate experiments.

---

**Table S1.** Description of the COVID-19 patient samples used in this study. All patients exhibited mild symptoms not requiring hospitalization except for patient 6 (indicated by an asterisk), who had moderate symptoms requiring non-invasive ventilation or a high flow O_2_ device.

| Participant ID | Day(s) post symptom onset (p.s.o.) | Age | Gender |
| --- | --- | --- | --- |
| 1 | 27, 72 | 47 | Female |
| 2 | 31, 60 | 43 | Female |
| 3 | 29, 71 | 65 | Male |
| 4 | 31, 67 | 29 | Male |
| 5 | 31, 63 | 48 | Female |
| 6* | 33, 76 | 64 | Female |
| 7 | 29, 74 | 22 | Male |
| 8 | 31, 67 | 31 | Female |
| 9 | 26, 55 | 56 | Male |
| 10 | 34, 67 | 28 | Female |
| 11 | 34, 62 | 30 | Male |
| 12 | 26, 58 | 36 | Female |
| 13 | 28, 66 | 65 | Male |
| 14 | 26, 64 | 65 | Female |
| 15 | 48, 77 | 52 | Female |
| 16 | 35, 69 | 52 | Male |
| 17 | 30, 57 | 36 | Male |
| 18 | 43, 73 | 29 | Male |

**Figure S1.**
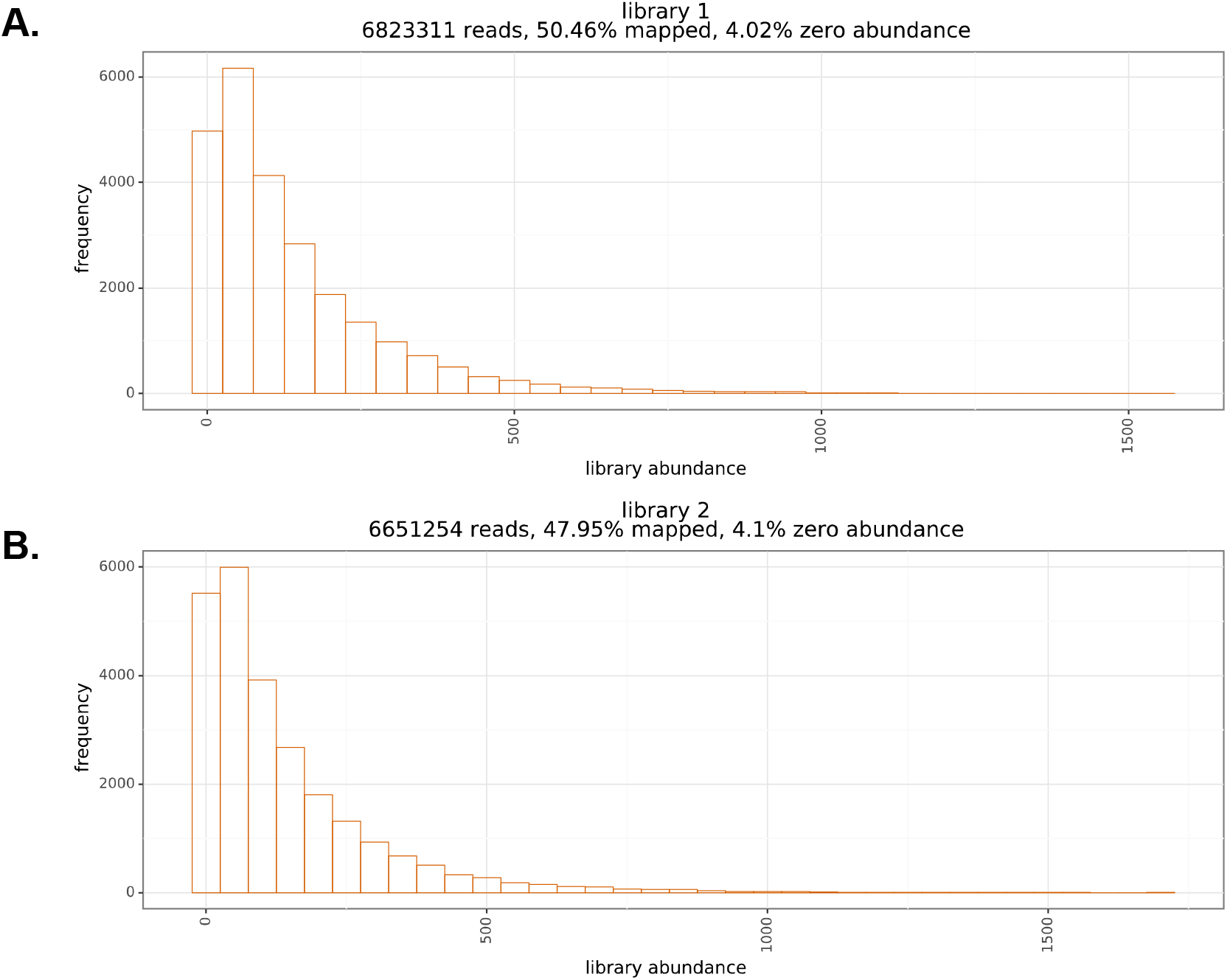
Distribution of sequenced peptides within biological replicate Spike Phage-DMS libraries. (A and B) Histogram showing the distribution of all sequenced peptides from a representative deep sequencing experiment for Spike Phage-DMS Library 1 (A) and Library 2 (B). Reads were stringently aligned to the reference library, allowing for 0 mismatches, and the proportion of unmapped reads is shown at the top. Additionally, the proportion of all non-sequenced peptides for each library is shown at the top.

**Figure S2.**
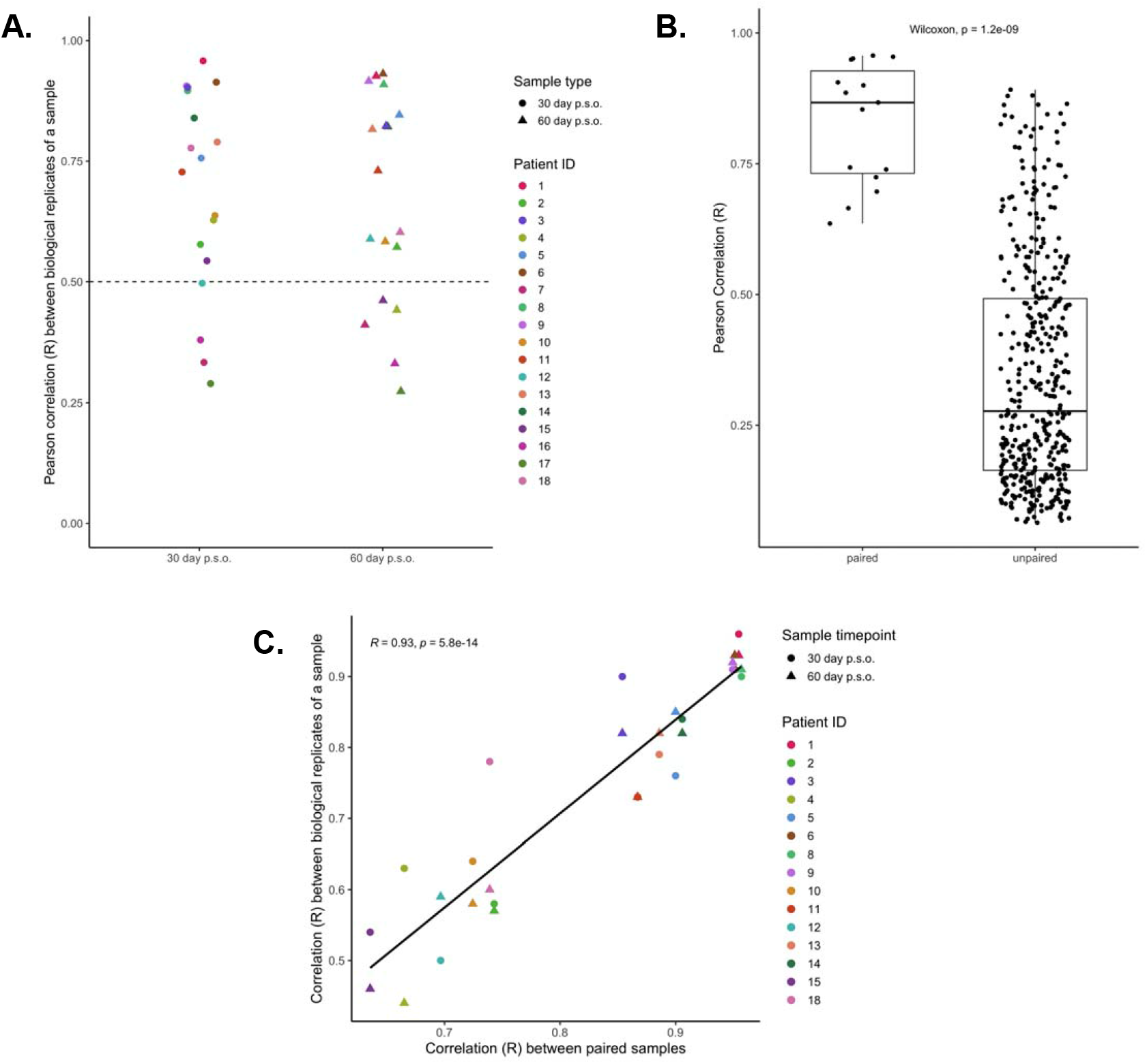
Reproducibility of peptide enrichment by plasma from COVID-19 patients. (A) Distribution of correlation values between peptide enrichment values for replicate experiments with samples from COVID-19 patients (Pearson’s correlation coefficient, R). Each color corresponds to a unique patient or volunteer, and the shape of each dot represents the type of sample. A dotted line at y = 0.5 represents the cutoff used to determine whether samples were kept in the analysis. (B) Boxplots showing the distribution of correlation values between patient samples that were paired between the day 30 and 60 p.s.o. timepoints (on the left) or samples that were randomly paired and compared (on the right). (C) Relationship between the biological replicate correlation for a sample and its correlation with its paired timepoint. Each color corresponds to a unique patient, and the shape of each dot represents the type of sample. Pearson’s correlation coefficient shown.

**Figure S3.**
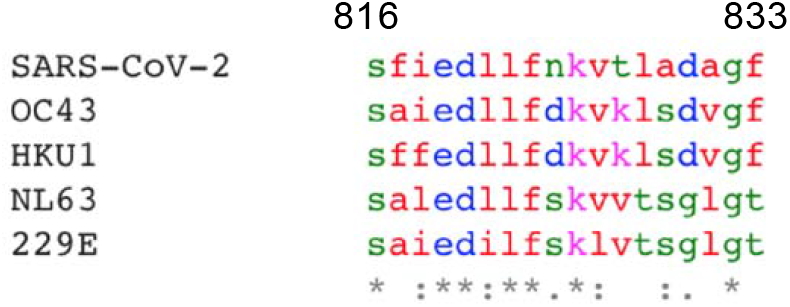
Multiple sequence alignment of the FP for SARS-CoV-2 and human endemic coronaviruses (OC43, HKU1, NL63, and 229E). Alignment was performed using Clustal Omega, and amino acids are colored according to physiochemical properties. GenBank accession numbers: YP_009724390.1, YP_009555241.1, YP_173238.1, YP_003767.1, and NP_073551.1, respectively.

**Figure S4.**
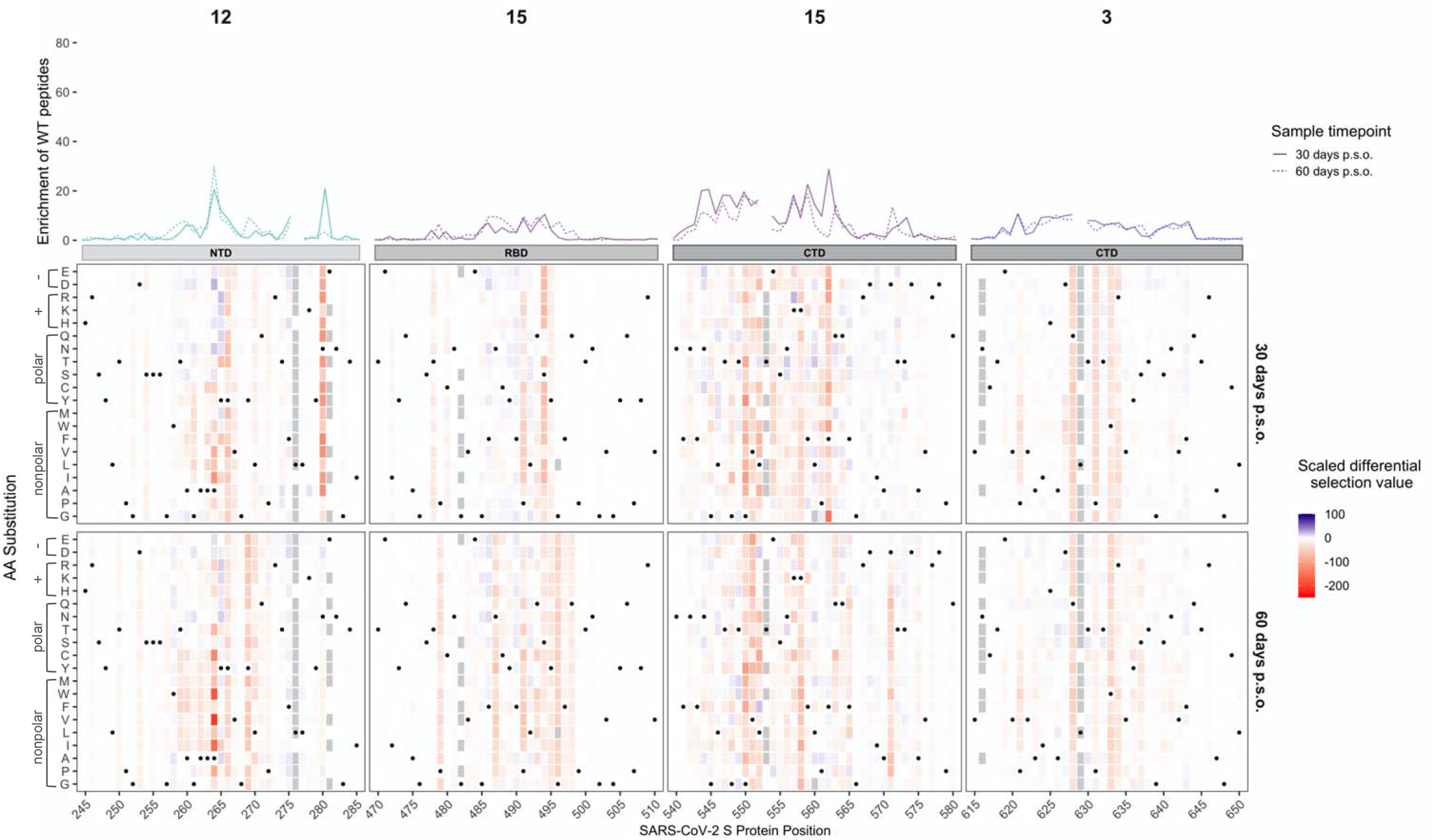
Effect of mutations on binding by COVID-19 patient plasma within various regions. Heatmaps depicting the effect of all mutations, as measured by scaled differential selection, at each site within the NTD, RBD, and CTD regions. Mutations enriched above the wildtype residue are colored blue and mutations depleted as compared to the wildtype residue are colored red. The wildtype residue is indicated with a black dot. Line plots showing the enrichment of wildtype peptides for each patient are shown above, with a solid line for patient samples taken at day 30 p.s.o. and a dashed line for patient samples taken at day 60 p.s.o. Peptides missing from the library are shown as grey boxes in the heatmaps and as breaks in the line plots.

## BIBLIOGRAPHY

Baum, A., Fulton, B.O., Wloga, E., Copin, R., Pascal, K.E., Russo, V., Giordano, S., Lanza, K., Negron, N., Ni, M., et al. (2020). Antibody cocktail to SARS-CoV-2 spike protein prevents rapid mutational escape seen with individual antibodies. Science 369, 1014–1018.

Belouzard, S., Chu, V.C., and Whittaker, G.R. (2009). Activation of the SARS coronavirus spike protein via sequential proteolytic cleavage at two distinct sites. Proc Natl Acad Sci U S A 106, 5871–5876.

Brouwer, P.J.M., Caniels, T.G., van der Straten, K., Snitselaar, J.L., Aldon, Y., Bangaru, S., Torres, J.L., Okba, N.M.A., Claireaux, M., Kerster, G., et al. (2020). Potent neutralizing antibodies from COVID-19 patients define multiple targets of vulnerability. Science 369, 643–650.

Di Tommaso, P., Chatzou, M., Floden, E.W., Barja, P.P., Palumbo, E., and Notredame, C. (2017). Nextflow enables reproducible computational workflows. Nat Biotechnol 35, 316–319.

Dingens, A.S., Arenz, D., Weight, H., Overbaugh, J., and Bloom, J.D. (2019). An Antigenic Atlas of HIV-1 Escape from Broadly Neutralizing Antibodies Distinguishes Functional and Structural Epitopes. Immunity 50, 520–532.e523.

Doud, M.B., Hensley, S.E., and Bloom, J.D. (2017). Complete mapping of viral escape from neutralizing antibodies. PLoS Pathog 13, e1006271.

Fan, X., Cao, D., Kong, L., and Zhang, X. (2020). Cryo-EM analysis of the post-fusion structure of the SARS-CoV spike glycoprotein. Nat Commun 11, 3618.

Garrett, M.E., Itell, H.L., Crawford, K.H.D., Basom, R., Bloom, J.D., and Overbaugh, J. (2020). Phage-DMS: A Comprehensive Method for Fine Mapping of Antibody Epitopes. iScience 23.

Gunn, B.M., Yu, W.H., Karim, M.M., Brannan, J.M., Herbert, A.S., Wec, A.Z., Halfmann, P.J., Fusco, M.L., Schendel, S.L., Gangavarapu, K., et al. (2018). A Role for Fc Function in Therapeutic Monoclonal Antibody-Mediated Protection against Ebola Virus. Cell Host Microbe 24, 221–233.e225.

Hassan, A.O., Case, J.B., Winkler, E.S., Thackray, L.B., Kafai, N.M., Bailey, A.L., McCune, B.T., Fox, J.M., Chen, R.E., Alsoussi, W.B., et al. (2020). A SARS-CoV-2 Infection Model in Mice Demonstrates Protection by Neutralizing Antibodies. Cell 182, 744–753.e744.

Haynes, B.F., Gilbert, P.B., McElrath, M.J., Zolla-Pazner, S., Tomaras, G.D., Alam, S.M., Evans, D.T., Montefiori, D.C., Karnasuta, C., Sutthent, R., et al. (2012). Immune-correlates analysis of an HIV-1 vaccine efficacy trial. N Engl J Med 366, 1275–1286.

Hilton, S.K.H., John; Black, Allison; North, Khrystyna; Dingens, Adam S.; Bedford, Trevor; Bloom, Jesse D. (2020). dms-view: Interactive visualization tool for deep mutational scanning data. Journal of Open Source Software 5.

Hoyer, S., and Hamman, J. (2017). xarray: N-D labeled Arrays and Datasets in Python. Journal of Open Research Software 5, 10.

Jiang, H.W., Li, Y., Zhang, H.N., Wang, W., Yang, X., Qi, H., Li, H., Men, D., Zhou, J., and Tao, S.C. (2020). SARS-CoV-2 proteome microarray for global profiling of COVID-19 specific IgG and IgM responses. Nat Commun 11, 3581.

Kleine-Weber, H., Elzayat, M.T., Wang, L., Graham, B.S., Müller, M.A., Drosten, C., Pöhlmann, S., and Hoffmann, M. (2019). Mutations in the Spike Protein of Middle East Respiratory Syndrome Coronavirus Transmitted in Korea Increase Resistance to Antibody-Mediated Neutralization. J Virol 93.

Korber, B., Fischer, W.M., Gnanakaran, S., Yoon, H., Theiler, J., Abfalterer, W., Hengartner, N., Giorgi, E.E., Bhattacharya, T., Foley, B., et al. (2020). Tracking Changes in SARS-CoV-2 Spike: Evidence that D614G Increases Infectivity of the COVID-19 Virus. Cell 182, 812–827.e819.

Langmead, B., Trapnell, C., Pop, M., and Salzberg, S.L. (2009). Ultrafast and memory-efficient alignment of short DNA sequences to the human genome. Genome Biol 10, R25.

Li, H., Handsaker, B., Wysoker, A., Fennell, T., Ruan, J., Homer, N., Marth, G., Abecasis, G., and Durbin, R. (2009). The Sequence Alignment/Map format and SAMtools. Bioinformatics 25, 2078–2079.

Li, Q., Wu, J., Nie, J., Zhang, L., Hao, H., Liu, S., Zhao, C., Zhang, Q., Liu, H., Nie, L., et al. (2020a). The Impact of Mutations in SARS-CoV-2 Spike on Viral Infectivity and Antigenicity. Cell 182, 1284–1294.e1289.

Li, Y., Lai, D.Y., Zhang, H.N., Jiang, H.W., Tian, X., Ma, M.L., Qi, H., Meng, Q.F., Guo, S.J., Wu, Y., et al. (2020b). Linear epitopes of SARS-CoV-2 spike protein elicit neutralizing antibodies in COVID-19 patients. Cell Mol Immunol.

Lip, K.M., Shen, S., Yang, X., Keng, C.T., Zhang, A., Oh, H.L., Li, Z.H., Hwang, L.A., Chou, C.F., Fielding, B.C., et al. (2006). Monoclonal antibodies targeting the HR2 domain and the region immediately upstream of the HR2 of the S protein neutralize in vitro infection of severe acute respiratory syndrome coronavirus. J Virol 80, 941–950.

Mabuka, J., Nduati, R., Odem-Davis, K., Peterson, D., and Overbaugh, J. (2012). HIV-specific antibodies capable of ADCC are common in breastmilk and are associated with reduced risk of transmission in women with high viral loads. PLoS Pathog 8, e1002739.

Milligan, C., Richardson, B.A., John-Stewart, G., Nduati, R., and Overbaugh, J. (2015). Passively acquired antibody-dependent cellular cytotoxicity (ADCC) activity in HIV-infected infants is associated with reduced mortality. Cell Host Microbe 17, 500–506.

Mohan, D., Wansley, D.L., Sie, B.M., Noon, M.S., Baer, A.N., Laserson, U., and Larman, H.B. (2018). PhIP-Seq characterization of serum antibodies using oligonucleotide-encoded peptidomes. Nat Protoc 13, 1958–1978.

Patel, J.S., Quates, C.J., Johnson, E.L., and Ytreberg, F.M. (2019). Expanding the watch list for potential Ebola virus antibody escape mutations. PLoS One 14, e0211093.

Pinto, D., Park, Y.J., Beltramello, M., Walls, A.C., Tortorici, M.A., Bianchi, S., Jaconi, S., Culap, K., Zatta, F., De Marco, A., et al. (2020). Cross-neutralization of SARS-CoV-2 by a human monoclonal SARS-CoV antibody. Nature 583, 290–295.

Poh, C.M., Carissimo, G., Wang, B., Amrun, S.N., Lee, C.Y., Chee, R.S., Fong, S.W., Yeo, N.K., Lee, W.H., Torres-Ruesta, A., et al. (2020). Two linear epitopes on the SARS-CoV-2 spike protein that elicit neutralising antibodies in COVID-19 patients. Nat Commun 11, 2806.

Robbiani, D.F., Gaebler, C., Muecksch, F., Lorenzi, J.C.C., Wang, Z., Cho, A., Agudelo, M., Barnes, C.O., Gazumyan, A., Finkin, S., et al. (2020). Convergent antibody responses to SARS-CoV-2 in convalescent individuals. Nature 584, 437–442.

Rogers, T.F., Zhao, F., Huang, D., Beutler, N., Burns, A., He, W.T., Limbo, O., Smith, C., Song, G., Woehl, J., et al. (2020). Isolation of potent SARS-CoV-2 neutralizing antibodies and protection from disease in a small animal model. Science 369, 956–963.

Saphire, E.O., Schendel, S.L., Gunn, B.M., Milligan, J.C., and Alter, G. (2018). Antibody-mediated protection against Ebola virus. Nat Immunol 19, 1169–1178.

Seydoux, E., Homad, L.J., MacCamy, A.J., Parks, K.R., Hurlburt, N.K., Jennewein, M.F., Akins, N.R., Stuart, A.B., Wan, Y.H., Feng, J., et al. (2020). Analysis of a SARS-CoV-2-Infected Individual Reveals Development of Potent Neutralizing Antibodies with Limited Somatic Mutation. Immunity 53, 98–105.e105.

Shang, J., Wan, Y., Luo, C., Ye, G., Geng, Q., Auerbach, A., and Li, F. (2020). Cell entry mechanisms of SARS-CoV-2. Proc Natl Acad Sci U S A 117, 11727–11734.

Shanmugaraj, B., Siriwattananon, K., Wangkanont, K., and Phoolcharoen, W. (2020). Perspectives on monoclonal antibody therapy as potential therapeutic intervention for Coronavirus disease-19 (COVID-19). Asian Pac J Allergy Immunol 38, 10–18.

Shrock, E., Fujimura, E., Kula, T., Timms, R.T., Lee, I.H., Leng, Y., Robinson, M.L., Sie, B.M., Li, M.Z., Chen, Y., et al. (2020). Viral epitope profiling of COVID-19 patients reveals cross-reactivity and correlates of severity. Science.

Sievers, F., Wilm, A., Dineen, D., Gibson, T.J., Karplus, K., Li, W., Lopez, R., McWilliam, H., Remmert, M., Söding, J., et al. (2011). Fast, scalable generation of high-quality protein multiple sequence alignments using Clustal Omega. Mol Syst Biol 7, 539.

Starr, T.N., Greaney, A.J., Hilton, S.K., Ellis, D., Crawford, K.H.D., Dingens, A.S., Navarro, M.J., Bowen, J.E., Tortorici, M.A., Walls, A.C., et al. (2020). Deep Mutational Scanning of SARS-CoV-2 Receptor Binding Domain Reveals Constraints on Folding and ACE2 Binding. Cell 182, 1295–1310.e1220.

Sui, J., Aird, D.R., Tamin, A., Murakami, A., Yan, M., Yammanuru, A., Jing, H., Kan, B., Liu, X., Zhu, Q., et al. (2008). Broadening of neutralization activity to directly block a dominant antibody-driven SARS-coronavirus evolution pathway. PLoS Pathog 4, e1000197.

van Rossum, G., and Drake, F.L. (2009). Python 3 Reference Manual (Scotts Valley, CA: CreateSpace).

Walls, A.C., Park, Y.J., Tortorici, M.A., Wall, A., McGuire, A.T., and Veesler, D. (2020). Structure, Function, and Antigenicity of the SARS-CoV-2 Spike Glycoprotein. Cell 181, 281–292.e286.

Walls, A.C., Tortorici, M.A., Snijder, J., Xiong, X., Bosch, B.J., Rey, F.A., and Veesler, D. (2017). Tectonic conformational changes of a coronavirus spike glycoprotein promote membrane fusion. Proc Natl Acad Sci U S A 114, 11157–11162.

Wan, J., Xing, S., Ding, L., Wang, Y., Gu, C., Wu, Y., Rong, B., Li, C., Wang, S., Chen, K., et al. (2020). Human-IgG-Neutralizing Monoclonal Antibodies Block the SARS-CoV-2 Infection. Cell Rep 32, 107918.

Wang, C., Li, W., Drabek, D., Okba, N.M.A., van Haperen, R., Osterhaus, A., van Kuppeveld, F.J.M., Haagmans, B.L., Grosveld, F., and Bosch, B.J. (2020). A human monoclonal antibody blocking SARS-CoV-2 infection. Nat Commun 11, 2251.

Wec, A.Z., Wrapp, D., Herbert, A.S., Maurer, D.P., Haslwanter, D., Sakharkar, M., Jangra, R.K., Dieterle, M.E., Lilov, A., Huang, D., et al. (2020). Broad neutralization of SARS-related viruses by human monoclonal antibodies. Science 369, 731–736.

Williams, K.L., Stumpf, M., Naiman, N.E., Ding, S., Garrett, M., Gobillot, T., Vézina, D., Dusenbury, K., Ramadoss, N.S., Basom, R., et al. (2019). Identification of HIV gp41-specific antibodies that mediate killing of infected cells. PLoS Pathog 15, e1007572.

Xia, S., Liu, M., Wang, C., Xu, W., Lan, Q., Feng, S., Qi, F., Bao, L., Du, L., Liu, S., et al. (2020). Inhibition of SARS-CoV-2 (previously 2019-nCoV) infection by a highly potent pan-coronavirus fusion inhibitor targeting its spike protein that harbors a high capacity to mediate membrane fusion. Cell Res 30, 343–355.

Yi, Z., Ling, Y., Zhang, X., Chen, J., Hu, K., Wang, Y., Song, W., Ying, T., Zhang, R., Lu, H., et al. (2020). Functional mapping of B-cell linear epitopes of SARS-CoV-2 in COVID-19 convalescent population. Emerg Microbes Infect 9, 1988–1996.

Zamecnik, C.R., Rajan, J.V., Yamauchi, K.A., Mann, S.A., Loudermilk, R.P., Sowa, G.M., Zorn, K.C., Alvarenga, B.D., Gaebler, C., Caskey, M., et al. (2020). ReScan, a Multiplex Diagnostic Pipeline, Pans Human Sera for SARS-CoV-2 Antigens. Cell Rep Med, 100123.

Zheng, Z., Monteil, V.M., Maurer-Stroh, S., Yew, C.W., Leong, C., Mohd-Ismail, N.K., Cheyyatraivendran Arularasu, S., Chow, V.T.K., Lin, R.T.P., Mirazimi, A., et al. (2020). Monoclonal antibodies for the S2 subunit of spike of SARS-CoV-1 cross-react with the newly-emerged SARS-CoV-2. Euro Surveill 25.

Zost, S.J., Gilchuk, P., Case, J.B., Binshtein, E., Chen, R.E., Nkolola, J.P., Schäfer, A., Reidy, J.X., Trivette, A., Nargi, R.S., et al. (2020). Potently neutralizing and protective human antibodies against SARS-CoV-2. Nature 584, 443–449.

Zulkower, V., and Rosser, S. (2020). DNA Chisel, a versatile sequence optimizer. Bioinformatics.

